# A Non-Negative Matrix Tri-Factorization based Method for Predicting Antitumor Drug Sensitivity

**DOI:** 10.1101/2021.12.03.471100

**Authors:** Sara Pidò, Carolina Testa, Pietro Pinoli

## Abstract

Large annotated cell line collections have been proven to enable the prediction of drug response in the preclinical setting. We present an enhancement of Non-Negative Matrix Tri-Factorization method, which allows the integration of different data types for the prediction of missing associations. To test our method we retrieved a dataset from CCLE, containing the connections among cell lines and drugs by means of their IC50 values. We performed two different kind of experiments: a) prediction of missing values in the matrix, b) prediction of the complete drug profile of a new cell line, demonstrating the validity of the method in both scenarios.

## 1 Scientific Background

Cancer is a highly complex disease due to the enormous level of both intra- and inter-tumor heterogeneity that often displays. Indeed, several tumors of the same organ may vary significantly in important tumor-associated attributes. This is the reason why patients with the same diagnosis can respond in different ways to the same therapy, and this represents the main obstacle to effective treatments [4].

The parameter most extensively used to characterize the response and sensitivity to a drug is the half-maximal inhibitory concentration (IC50), i.e., the concentration needed to inhibit the 50% of the targeted biological process or component [5]. In particular, in the field of anticancer therapies, it represents the concentration of drugs needed to kill half of the cells. Since experimental approaches for the estimation of IC50 values are costly and time-consuming, researchers are increasingly putting efforts into developing computer-based methods for predicting the responsiveness of a patient to a drug. Therefore, in the context of precision medicine, the prediction of drug sensitivity became of fundamental relevance. Being able to determine a priori to which drugs a patient, with its genomic features, is sensitive or resistant would save a lot of precious time and improve the efficiency of the therapy.

In this scenario, to predict the sensitivity of a cell line to a drug, we used a network-based approach based on Non-negative Matrix Tri-Factorization (NMTF), an algorithm designed to factorize an input association matrix in three matrices of non-negative elements in order to predict missing associations. One of the main advantages of NMTF is the possibility to expand the network integrating several information; indeed, it allows to start from a bipartite graph, containing the connection we want to predict, and to add further related data to build a multipartite graph which decomposes each association matrix [2]. The NMTF approach demonstrates to have elevated performances in both finding new indications for approved drugs and new synergistic drug pairs, in particular when including several heterogeneous data types [2, 6]. The main focus of this work is to adapt the model to predict the sensitivity of a cancer cell line to a set of antitumor drugs integrating the associations between cell lines and drugs with tissue and gene expression-related data. The prediction of drug response based on genetic features could speed the emergence of ‘personalized’ therapeutic regimens.

## 2 Data and Methods

### 2.1 Dataset

For our experiments, we used a dataset retrieved from the Cancer Cell Line Encyclopedia (CCLE) [1] which comprises the association among 1065 cell lines and 266 antitumor drugs, measured in terms of IC50. The IC50 is representative of the response of a cell line to a drug: we consider a cell line to be sensitive (resistant) to a drug if the IC50 value is lower (higher) than the median of IC50 values of that drug considering all the cell lines. For 647 of those cell lines, further information is available; in particular, we considered tissues of origin and the gene expression quantification by RNA-seq experiments.

### 2.2 Model

In order to integrate all the available information, we modeled the set of cell lines *C*, the set of drugs *D*, the set of tissues *T* and the set of genes *G* as the multipartite network in Fig. 1, where each cell line is connected to the drugs to which it is sensitive, the tissue of origin and a set of genes, with the weight of the edge representing the expression of the gene in the cell line.

**Figure 1:**
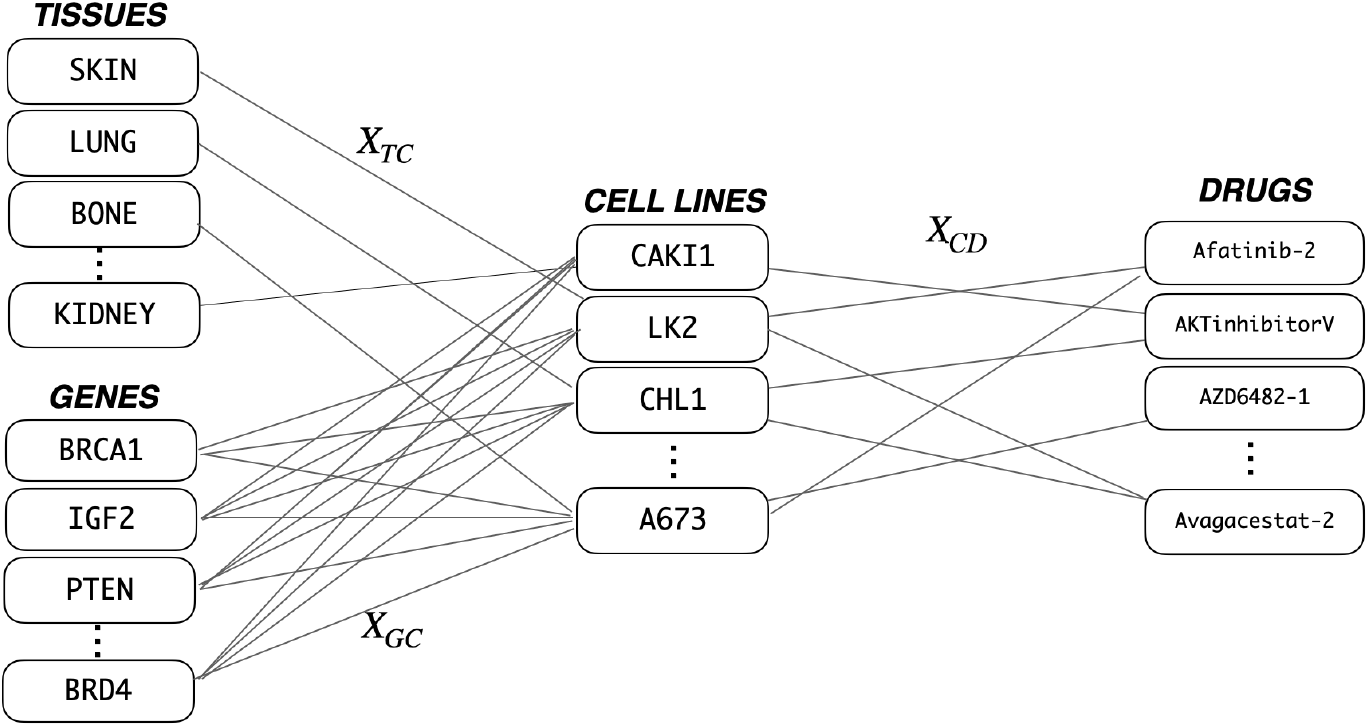
Multipartite graph connecting Tissues, Gene expression, Cell lines and Drugs. The three association matrices of the graph are also indicated.

Such network is equivalent to the set of its association matrices: a binary matrix *X*_*CD*_ connecting cell lines to drugs, a binary matrix *X*_*TC*_ connecting cell lines to tissues, and a real matrix *X*_*GC*_ connecting cell lines to genes. We build the three matrices as follows:

- we represent the *IC*50 data as a matrix 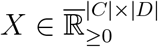, being 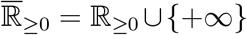, such that *X*[*i, j*] indicates the IC50 value of the j-*th* drug on the i-*th* cell line if a measure is available, or +∞ otherwise. We transform *X* into the binary matrix *X*_*CD*_ ∈ {0, 1}^|*C* |*×*|*D*|,^such that:

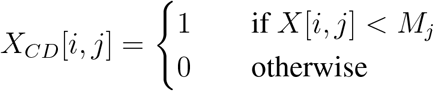

- where *M*_*j*_ is the median of the IC50 values for the drug *j* different from + ∞ In other words, an element of *X*_*CD*_[*i, j*] is equal to 1 only if the i-*th* cell line is sensitive to the the j-*th* drug;
- we build the matrix *X*_*TC*_ *∈* {0, 1}^|*T* |*×*|*C*|^ that connects cell lines to tissues as:

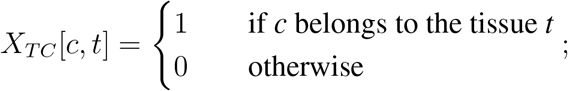

we consider the matrix 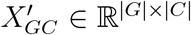, where *G* is a set of genes and 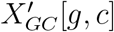 represents the *RPKM*, i.e., *reads per kilobase of transcript, per million mapped reads*, measured for the given gene *g* in the specific cell line *c*, by a RNAseq experiment. Finally, we keep into consideration *X*_*GC*_ ∈ *ℝ*^1000*×*|*C*|^ as the matrix containing only the 1000 genes with the highest variability across the cell lines.

### 2.3 Method

Let’s consider a multipartite graph 𝒢; for the purpose of this work, we can represent the graph as a set of association matrices, i.e., 𝒢 = {*X*_*IJ*_}, such that each association matrix 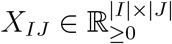 connects nodes of a set *I* to nodes of a set *J*.

We can apply the NMTF method to factorize each association matrix *X*_*IJ*_ into three matrices:

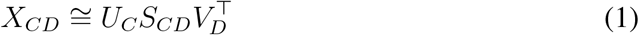

where 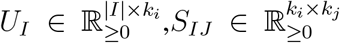, and 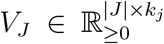 with *k*_*i*_, *k*_*j*_ *∈ ℕ* and *k*_*i*_ *<* |*I*| *k*_*j*_ *<*|*J*| The Parameters *k*_*i*_ and *k*_*j*_ are the factorization ranks of NMTF and describe the number of hidden vectors into which we want to represent the *X*_*IJ*_ association matrix.

Furthermore, the following constraint has to hold:

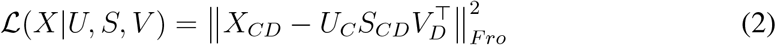

The factorization matrices are computed so as to minimize the objective function based on the Frobenius norm:

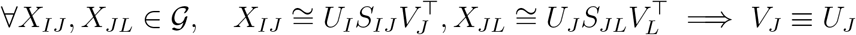

where Θ represents the set of all the factorization matrices.

A minimum of the objective function can be computed algorithmically by (a) initializing the factorization matrices with random positive numbers and (b) iteratively applying the following multiplicative update rules until convergence (e.g.,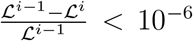) [3]:

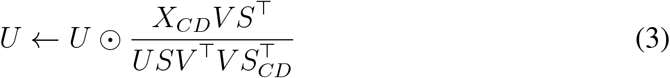

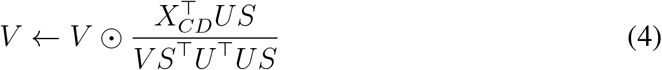

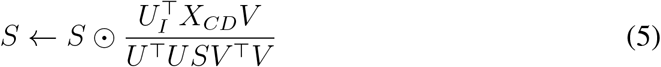

where ⊙ and 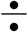 stand for Hadamard element-wise multiplication and division, respectively.

### 2.4 Prediction of Novel Associations

The prediction of novel associations between two sets of nodes can be interpreted as a matrix completion task. The NMTF method is applied in order to predict novel links between two classes of nodes. In particular, we focus on the associations between cell lines and drugs. After that

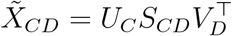

has been computed, we apply a threshold *τ*, typically 0 *< τ <* 1, and we consider that the i-*th* cell line is associated with the j-*th* drug if the predicted value 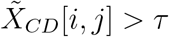.

### 2.5 Prediction of Drug Profile for New Cell Line

Another scenario is when a novel cell line is included in the network. In this situation, while we know the genetic feature of the cell line and its tissue of origin, we do not have information about the drugs to which it is sensitive.

We here propose a slight modification of the NMTF multiplicative update rules, in order to being able to predict the complete drug profile for the novel cell line. Since we have no correct information in the matrix we aim to reconstruct for the novel cell line, we do not consider the influence of *X*_*CD*_ during the update of *U*_*C*_ matrix. Thus, the new update rule (for our network) is:

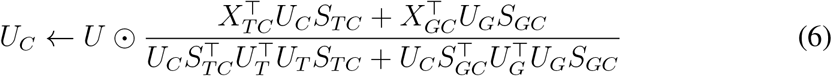

## 3 Results

Here, we report the results of different trials that we performed on the dataset illustrated in Section 2.1. In particular, we apply NMTF method, illustrated in Section 2.3, for two different tasks: the prediction of novel cell line-drug association and the prediction of the drug profile for a new cell line. We evaluate our results using the AUROC (i.e., area under the receiver operating characteristic curve) and the comparison between the actual IC50 values of pairs predicted sensitive 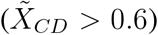 and predicted resistant 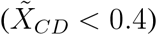.

### 3.1 Prediction of Novel Associations

In order to validate the model, we apply a mask that covers randomly the 5% of the association matrix *X*_*CD*_. We run the method on the single matrix *X*_*CD*_ without passing other information and we computed the evaluation metrics. The method performs well, indeed the AUROC is 0.88846 and according to the Welch test, the predicted sensitive and resistant associations are significantly different (p-val = 9.959611e-14). Results are shown in Fig. 2a, 2b.

**Figure 2:**
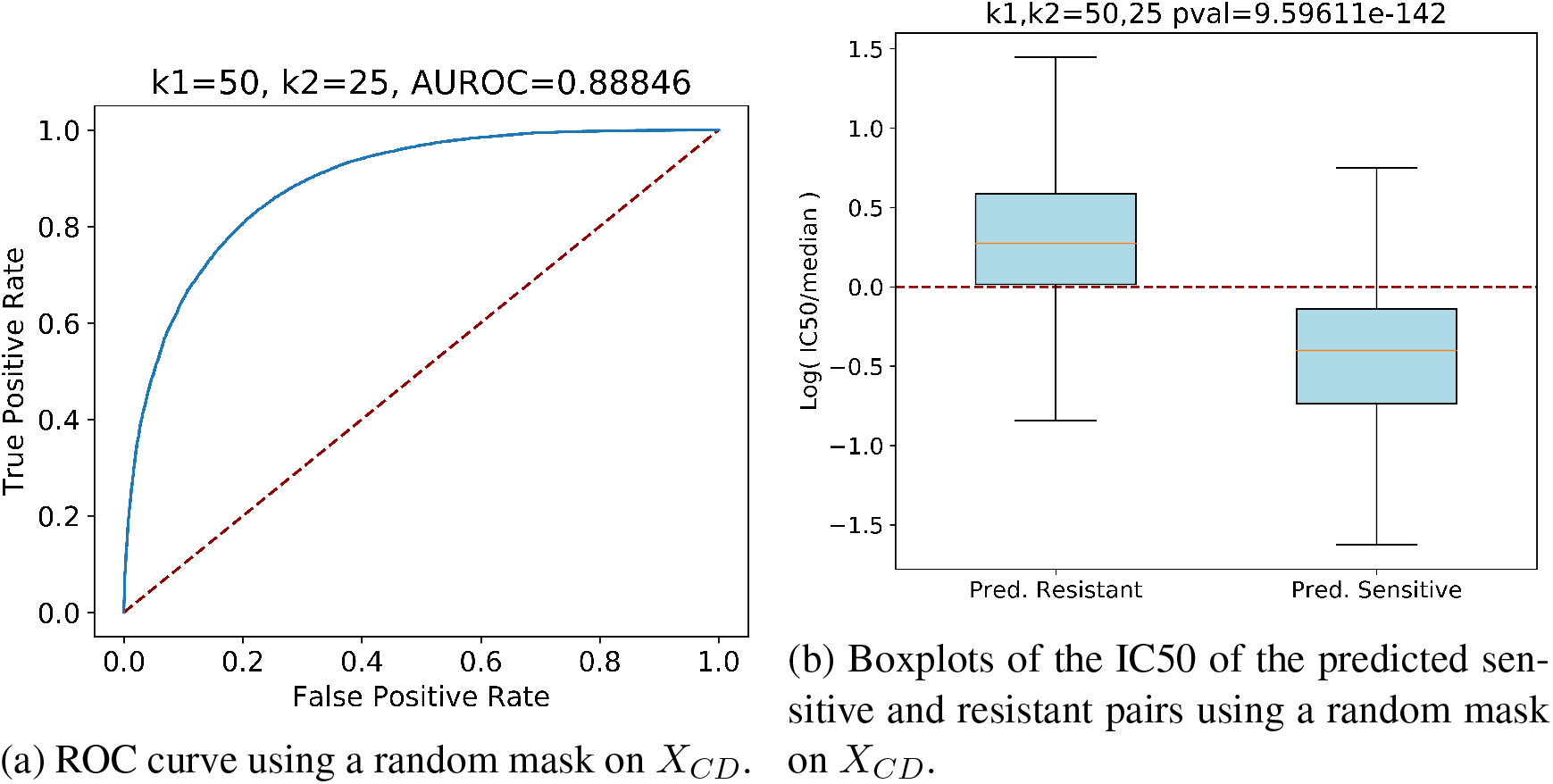
Performances using a random mask on *X*_*CD*_ with k1=50 and k2=25, where k1 and k2 are the factorization ranks of the NMTF

### 3.2 Prediction of Drug Profile for New Cell Line

In this case, we apply a mask on a single row of the matrix *X*_*CD*_ in order to simulate the addition of a novel cell line.

Considering *only X*_*CD*_ matrix does not provide meaningful results, as shown in Fig. 3a, 3b. As expected, without any additional information, the AUROC is 0.50449, and the two classes are not different. This result proves that it is impossible to predict a complete drug profile for a novel cell line without considering other data.

**Figure 3:**
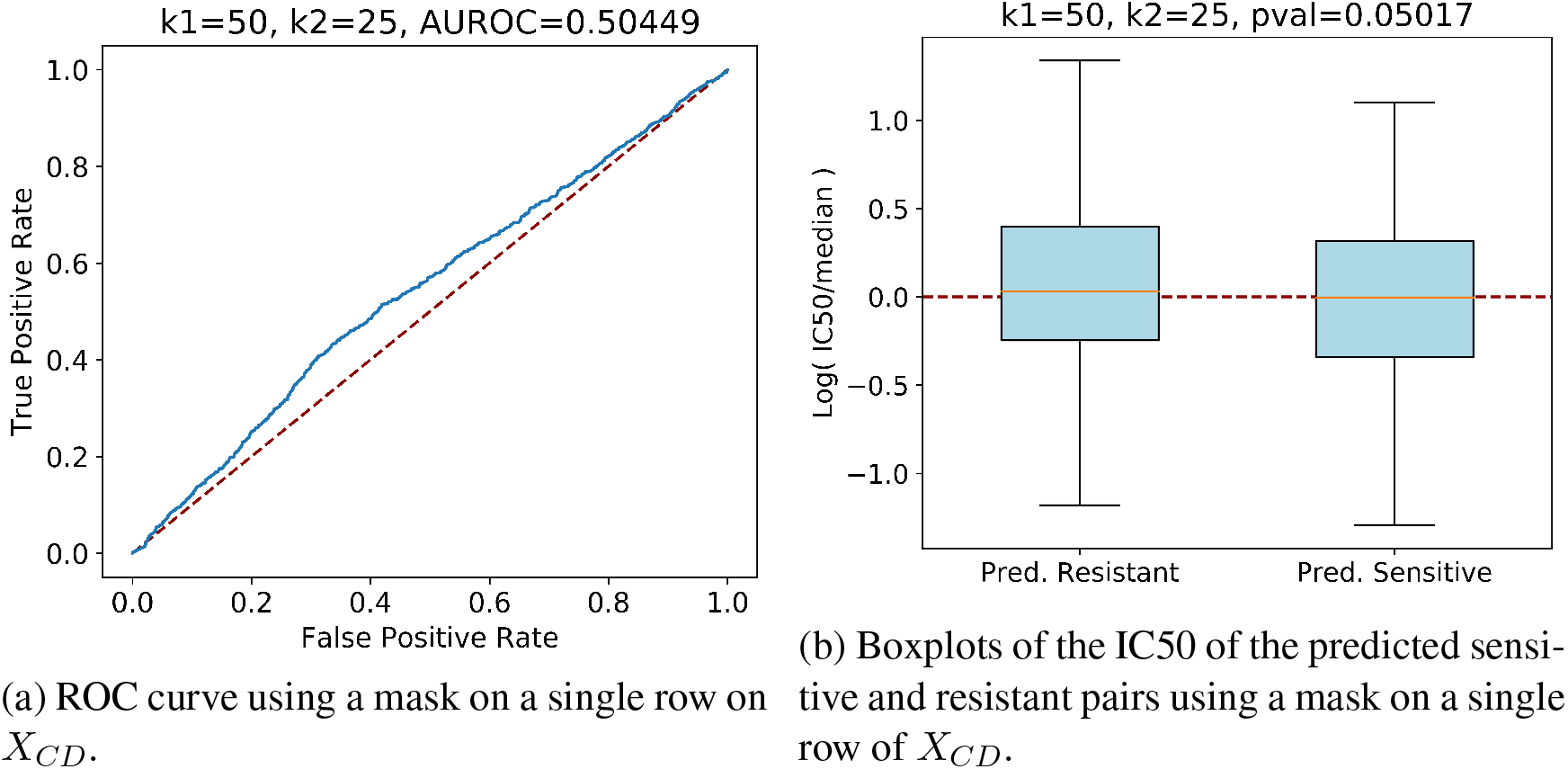
Performances using a mask on a single row of *X*_*CD*_ with k1=50 and k2=25, where k1 and k2 are the factorization ranks of NMTF.

Thus, we test the method by also adding the *X*_*T C*_ matrix alone, *X*_*GC*_ matrix alone as well as the two together.

The AUROCs in Fig. 4a proves that adding information increases the performances of the predictor. Including the tissue of origin, the method is able to reach an AU-ROC=0.60666. If also gene expressions are added to the model, we observe a significant improvement (AUROC=0.71094). Finally, when both gene expressions and tissues of origin are considered, and the AUROC increases to 0.72885. In Fig. 4b the comparison between predicted resistant and sensitive drugs, when all the information is used, is shown; the Welch test confirms the difference in the distribution of the two classes (p-val = 1.40798e-34), with the IC50 of the predicted sensitive drugs clearly below the median.

**Figure 4:**
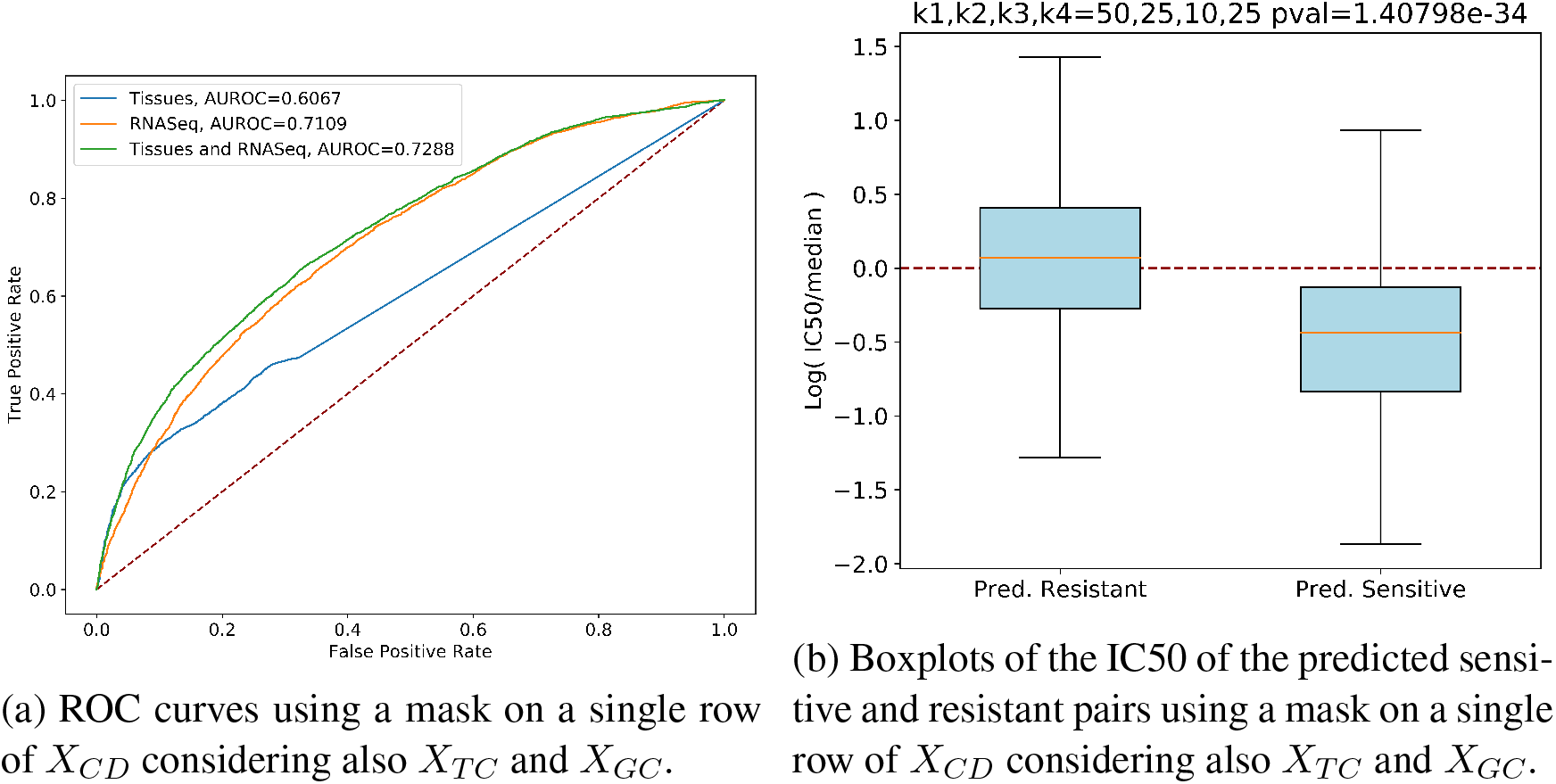
Performances using a mask on a single row of *X*_*CD*_ considering also *X*_*TC*_ and *X*_*GC*_ with k1=50, k2=25, k3=10, k4=25, where k1, k2, k3, k4 are the factorization ranks of NMTF.

## 4 Conclusions

In our work, we demonstrated that predicting the sensitivity of a specific drug for a given cell line for which many IC50 experiments are available is a rather easy task. In our experiments, using plain NMTF method without additional information for this task allows to reach high performances (AUROC = 0.88846). On the contrary, predicting drug sensitivity profile for a novel cell line is a complex task: indeed, NMTF method without other data scores as bad as a random predictor.

To overcome this limitation, we proposed a two-fold solution: (a) we developed an improved version of NMTF algorithm, which generates predictions taking into account only meaningful information, and (b) we integrated other information, namely the tissues of origin and the gene expressions of the corpus of cell lines.

When all the available data are provided, the proposed method shows much better performances: the resulted AUROC is equal to 0.72885. In particular, we observed how gene expressions is fundamental in order to improve the prediction.

As future development we would like to enlarge the network to further improve the performance. In particular, we would like to test if generating tissue-specific networks may provide a positive influence to the prediction. Moreover, we want to implement a regression method in order to being able to predict also the weight of the connection, i.e., the IC50 value. This could certainly help to find more rapidly the right therapy for the patient, saving time and improving its treatment.

